# Beyond Structure and Affinity: Context-Dependent Signals for de novo Binder Success

**DOI:** 10.64898/2026.04.13.718094

**Authors:** Çağlar Bozkurt

## Abstract

De novo protein binder design has advanced rapidly, yet most designs fail experimentally and current structure- and affinity-centred evaluation does not reliably predict which candidates will succeed. Here we show that biology-informed sequence features, derived from models trained on natural proteins, identify transferable and context-dependent associations with binder expression and binding that are not captured by structural scoring alone.

We re-analysed two public benchmarks—the Bits to Binders CAR-T CD20 competition (11,984 designs; expression, proliferation, and T cell function gates) and the Adaptyv EGFR competition (603 designs; expression and binding affinity)—using five biology-informed ML models predicting disorder, amyloidogenicity, topology, PTM sites, and protein classification. Every feature was tested at each gate with FDR-corrected statistics.

We identify three layers of signal. *Transferable*: lower aggregation propensity is the most robust cross-benchmark signal; PTM-site density recurs univariately but is partly length-confounded in EGFR. *Architecture-dependent*: topology, disorder, and disulfide-related descriptors are significant in both datasets but flip direction, consistent with the different requirements of CAR extracellular domains versus standalone binders. *Context-specific*: phosphorylation-related associations with CAR-T depletion and low-disorder dominance in EGFR binding are tied to individual assay or format contexts. In the CAR-T benchmark, stacking biology-informed filters raises the enrichment hit rate from 13.8% to 38.6% (2.8× lift) after controlling for known sequence-level predictors.

These results suggest that pre-synthesis screening of de novo binders may benefit from being multi-gate and context-aware, using biology-informed sequence descriptors not only to rank candidates but also to help flag likely failure modes earlier and reduce wasted synthesis and testing.

## Introduction

### Binder Evaluation Remains Overfocused on Structure and Affinity

Computational protein binder design has matured to the point where generative models can propose thousands of candidate sequences for a given target [3, 4]. Evaluation of these candidates typically relies on structural plausibility metrics—predicted local distance difference test (pLDDT) scores, interface predicted aligned error (ipTM), Rosetta interface energies—and on binding affinity proxies such as docking scores or ProteinMPNN likelihoods [5, 4]. These metrics assess whether a designed sequence is likely to fold into a stable structure and make productive contacts with the target.

However, recent large-scale experimental benchmarks have revealed a disconcerting gap between *in silico* confidence and *in vivo* function. In the Bits to Binders competition, which tested 12,000 AI-designed CAR-T binder domains in human T cells, structural confidence scores (pLDDT, ipTM) and deep-learning-based likelihoods (ProteinMPNN, ESM-2 PLL) had limited predictive power for functional success [1], while simple sequence-level heuristics—such as K+E amino acid content and cysteine count— outperformed complex scoring functions. A similar pattern was observed in the Adaptyv EGFR binder competition, where hit rates remained modest despite sophisticated generative pipelines [2].

### The Missing Layer: Expression, Compatibility, and Context-Dependent Biology

These observations suggest that something beyond structure and affinity influences whether a designed binder succeeds in practice. A binder must not only fold and dock correctly; it must also:

- be expressible in the relevant host system and avoid aggregation or misfolding during biosynthesis;
- be compatible with the architectural context in which it operates (e.g., membrane-anchored versus standalone);
- produce productive downstream biology, not just target binding.

These are fundamentally *biological compatibility* constraints. They depend on the sequence’s relationship to the cellular machinery that synthesises, folds, traffics, and deploys it—information that is not captured by structure prediction or docking scores alone.

### Research Question

We ask: can biology-informed sequence features—derived from models trained to predict biological function on natural proteins—identify signals associated with binder success and experimental failure that complement structure- and affinity-centred scoring, and that could help flag failure risks before candidates reach the laboratory? And do these signals generalise across distinct binder architectures and assay settings, or are they context-dependent? More specifically, we ask whether such descriptors can help distinguish likely failure modes across experimental gates and deployment contexts, rather than only separating broadly successful from unsuccessful candidates.

### Public Benchmarks and Study Design

To answer these questions, we re-analysed two large public binder-design competitions that impose genuinely different design constraints. The CAR-T benchmark was analysed first to establish multi-gate associations; the EGFR benchmark was then used to test whether those associations generalise or change direction in a different architectural context:

#### 1. Bits to Binders CAR-T CD20 benchmark [1]

11,984 AI-designed 80-amino-acid binders targeting human CD20—a four-pass transmembrane protein with small, membrane-proximal extracellular loops. Binders were tested as the extracellular domain of a CAR28z receptor [7] in primary human T cells, through a multi-gate evaluation from DNA synthesis through expression, CD20-specific proliferation enrichment, and CD20-specific depletion. In this context, a binder must not only engage its target but also express on a T cell surface, fold within the CAR construct, and support productive downstream signalling.

#### 2. Adaptyv EGFR benchmark [2]

603 binder designs of variable length (13– 250 aa) targeting human EGFR—a single-pass receptor tyrosine kinase with a large, accessible extracellular domain. Binders were tested as standalone proteins via cell-free expression and bio-layer interferometry for binding affinity. Here, a binder must fold independently into a compact, stable structure and engage a large target surface without the constraints of a membrane receptor construct.

**Figure 1:**
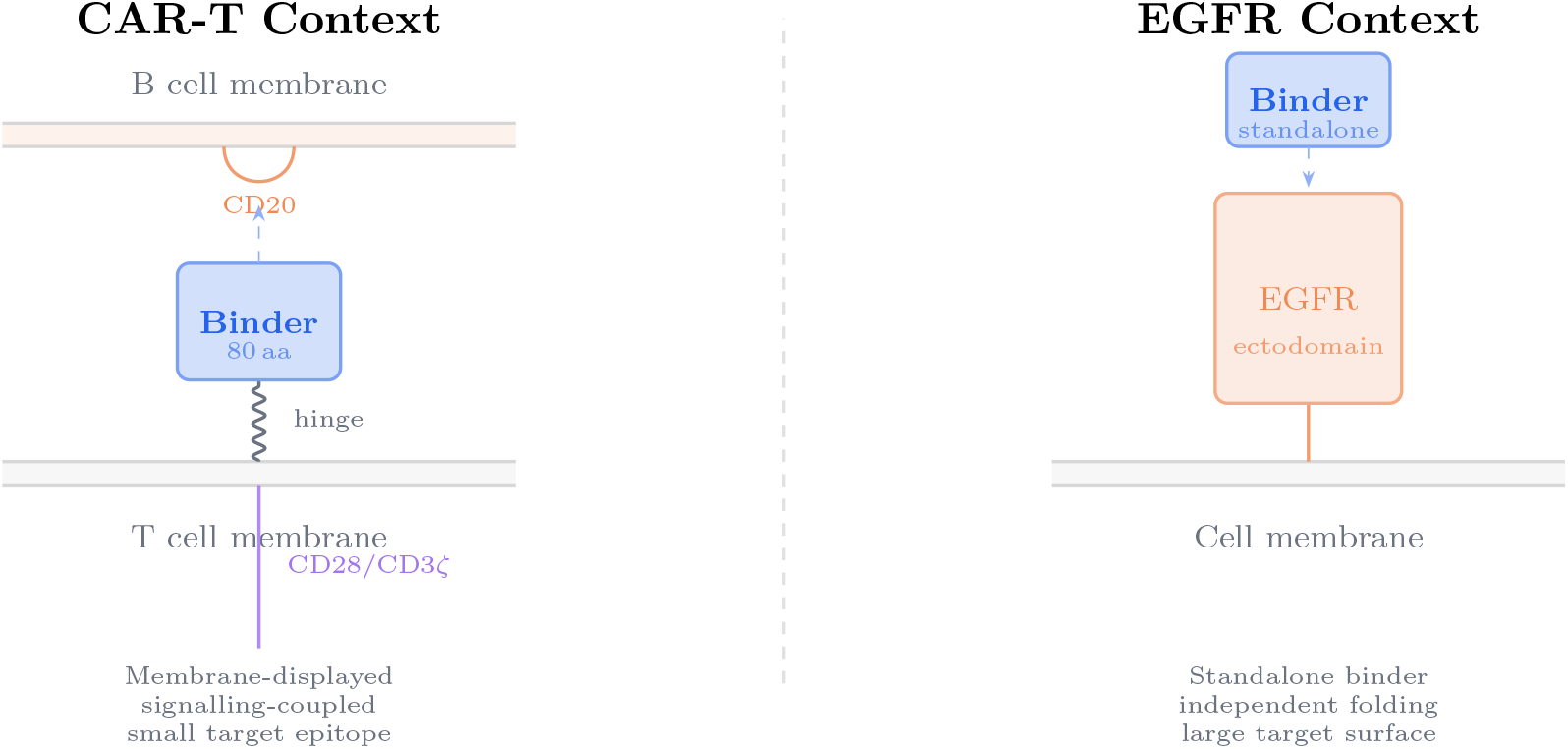
Architectural contexts impose different design constraints. Left: in the CAR-T benchmark, the binder operates as a membrane-displayed extracellular domain coupled to signalling machinery, engaging a small, membrane-proximal target. Right: in the EGFR benchmark, the binder must fold independently and engage a large extracellular target surface. These different biological and deployment contexts impose different structural and compatibility requirements on the binder sequence.

For every sequence in both datasets, we generated predictions from five biology-informed ML models and tested each feature at each experimental gate with FDR-corrected statistics, then compared results across datasets.

**Figure 2:**
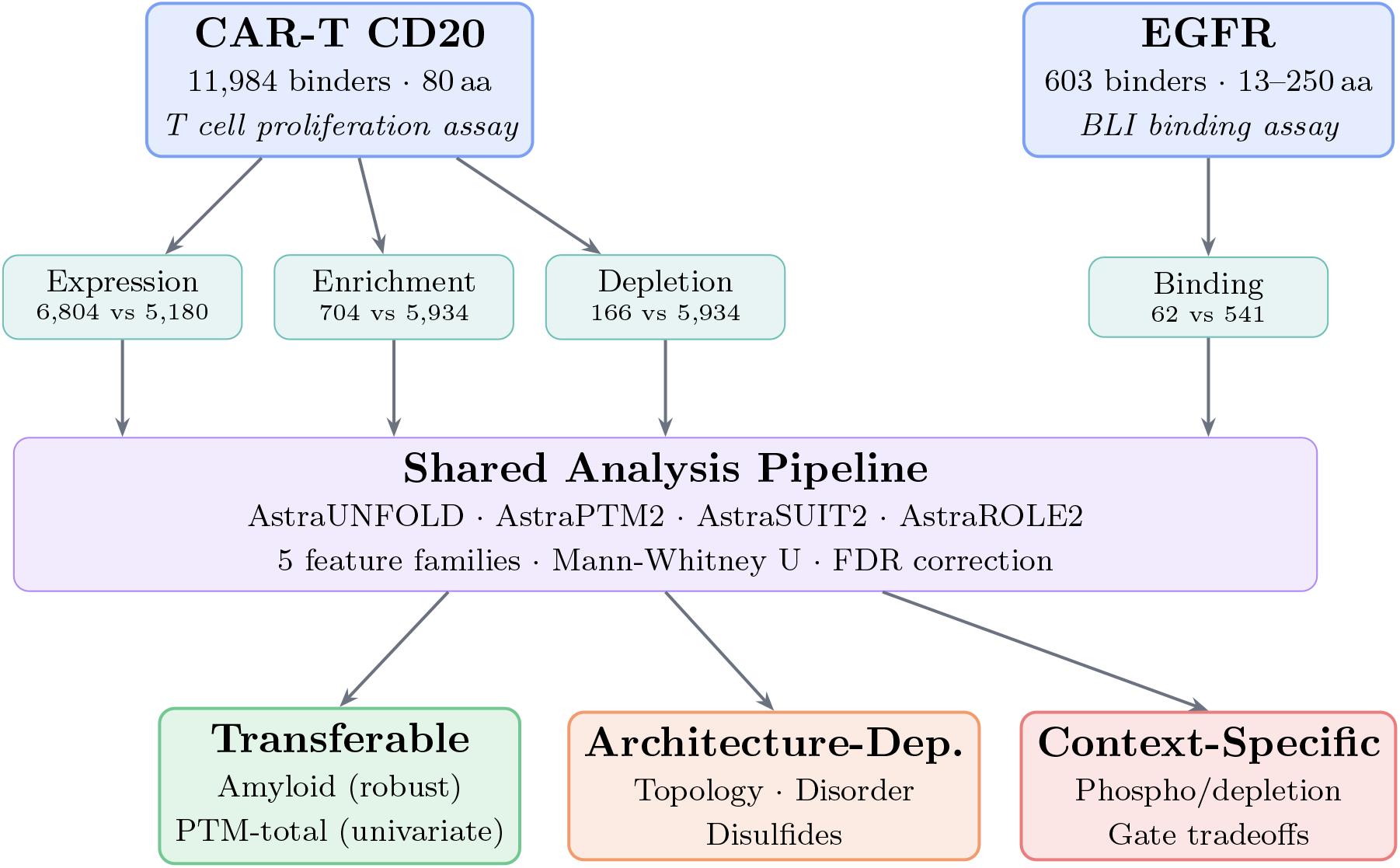
Study design. Two public binder-design competition datasets were analysed with a shared biology-informed feature pipeline. Features were tested at each experimental gate and classified as transferable, architecture-dependent, or context-specific based on cross-benchmark comparison.

## Methods

### Datasets

#### CAR-T CD20 Benchmark

The Bits to Binders competition [1] collected 12,000 de novo binder designs from 28 teams across 42 countries. Each design was an 80-amino-acid protein intended to bind the extracellular domain of human CD20 and serve as the extracellular domain of a second-generation CAR28z receptor.

Designs were codon-optimised, synthesised, cloned into a CAR backbone, and tested in primary human T cells through a series of experimental gates: DNA synthesis QC; NGS recovery after a pooled T cell proliferation assay (expression proxy); CD20-specific enrichment and depletion measured by edgeR differential analysis of barcode read counts; and individual functional assays (cytotoxicity, cytokine production, proliferation, expansion) for the top 10 candidates.

From this data, we defined four testable gate comparisons:

- **Expression** (recovery as a proxy for successful protein expression and T cell viability): recovered (n=6,804) vs. not recovered (n=5,180)
- **Enrichment** (productive CD20 engagement driving T cell proliferation): enriched (n=704) vs. neutral (n=5,934)
- **Depletion** (harmful or dysfunctional engagement causing T cell dropout): depleted (n=166) vs. neutral (n=5,934)
- **Enriched vs. Depleted**: opposite functional outcomes (n=704 vs. 166)

A controlled subset (cysteine-free sequences with K+E amino acid fraction *<*30%; n=4,020) was used to isolate signals independent of the paper’s known predictors.

#### EGFR Benchmark

The Adaptyv EGFR Binder Design Competition [2] ran two rounds with 159 participants across both rounds. Round 1 released 202 tested designs (*<*200 aa; BindCraft was the top-performing method in this round). Round 2 released 401 tested designs (*<*250 aa), selected from a larger submission pool by a weighted ranking of ESM2-PLL and iPAE+ipTM scores. Both rounds used cell-free protein synthesis for expression and bio-layer interferometry (BLI) for binding affinity measurement.

Round 1 and Round 2 sequences are fully distinct (zero overlap), giving 603 unique sequences in total. Gate comparisons:

- **Binding**: binders (n=62) vs. non-binders (n=541)
- **Expression**: expressed (n=379) vs. not expressed (n=22; Round 2 only; treated as exploratory due to small *n*)
- **Binding strength**: strong+medium (n=30) vs. weak+none (n=349; Round 2 only)

Sequence lengths ranged from 13 to 250 aa (mean 105), spanning miniproteins, nanobodies, peptides, and scFv fragments.

### Biology-Informed Feature Generation

For every sequence in both datasets, we generated predictions from five models in the Orbion Astra suite:

- **AstraUNFOLD**: per-residue predictions for intrinsic disorder, amyloidogenicity, and membrane topology (seven topology labels).
- **AstraPTM2** [12]: residue-level post-translational modification site predictions for 40 PTM types, combining standard and experimentally calibrated prediction modes.
- **AstraPTM-Mini**: protein-level binary presence/absence predictions for PTM types.
- **AstraSUIT2** [13]: multi-task protein classification covering taxonomic domain, host organism, membrane type, subcellular localisation, cofactor binding, and quaternary structure.
- **AstraROLE2** [13]: multi-task functional annotation covering EC numbers, Gene Ontology terms, protein categories, and metabolic pathway memberships.

All models use ESM-2 embeddings [6] and physicochemical descriptors as input. They were trained on natural protein datasets and are not used here as literal annotators of synthetic binder sequences. Instead, we treat them as *biology-informed sequence descriptors*: their outputs encode patterns learned from natural proteins—such as local charge distribution, hydrophobic burial, and surface exposure character—that may be informative about sequence compatibility even when applied outside their original training domain. A “predicted PTM site,” for example, should be read as a sequence neighbourhood that *resembles* a known modification context, not as a claim that the binder will be post-translationally modified. Similarly, “topology-like” outputs reflect sequence character associated with membrane topology in natural proteins, not literal topology assignments for synthetic constructs. This framing applies throughout the paper: when we refer to PTM-site density, topology character, or other model outputs, we mean the sequence-descriptor value, not a confirmed biochemical annotation.

In particular, PTM-related predictions applied to binders tested in cell-free expression systems (EGFR benchmark) should be interpreted primarily as sequence-character proxies rather than predictions of actual post-translational modification events. This duality—encoding biological regularities while not necessarily predicting actual biochemistry for synthetic sequences—is discussed further in the Discussion.

### Feature Grouping and Regional Definitions

Extracted features were organised into five groups:

1. **Aggregation/amyloid**: mean probability and fraction of residues predicted as amyloidogenic.
2. **Disorder**: mean probability and fraction of residues predicted as disordered.
3. **Topology-like character**: fraction of residues with “Outside”, “Inside”, or “Membrane” topology predictions. We use the term “topology-like” because these outputs reflect sequence character associated with membrane topology in natural proteins, not literal topology assignments for synthetic binders.
4. **PTM-related features**: total predicted PTM site counts and per-type counts for major modification types.
5. **Model label probabilities**: individual label probabilities from every AstraSUIT2 and AstraROLE2 prediction task.

For regional analysis, features were computed for whole sequences and for three subregions. In the CAR-T dataset (fixed 80-aa length), regions were defined as N-terminal (positions 1–20), core (21–60), and C-terminal (61–80). In the EGFR dataset (variable length), regions were defined proportionally: first 25%, middle 50%, and last 25% of residues.

### Statistical Analysis

For each gate comparison in each dataset, every continuous feature was tested using a two-sided Mann-Whitney U test, and every categorical feature using a chi-squared test. Effect sizes were computed as rank-biserial correlation *r* (for Mann-Whitney) and Cramér’s *V* (for chi-squared). Benjamini-Hochberg FDR correction [9] was applied across all tests within each gate.

In the CAR-T dataset, all comparisons were run twice: on the full dataset (“raw”) and on a controlled subset filtered to cysteine-free sequences with K+E fraction *<*30% (“controlled”), to isolate signals independent of the original paper’s known predictors. Significance thresholds: FDR *<* 0.05 for statistical significance; |*r*| *>* 0.15 for medium effect size; |*r*| *>* 0.30 for large effect size.

### Cross-Benchmark Comparison Framework

To compare results across datasets, we classified features into three categories based on a post-hoc synthesis of the per-gate, FDR-corrected univariate results. This taxonomy is a descriptive organising framework, not a separately corrected inferential layer; no additional multiple-testing correction was applied to the cross-benchmark classification itself.

- **Transferable**: same directional association with binding success, statistically significant in both datasets.
- **Architecture-dependent**: statistically significant in both datasets but with opposite directional association, consistent with the different structural requirements of each binder architecture.
- **Context-specific**: significant in only one dataset or at only one experimental gate, reflecting assay- or format-specific biology.

## Results

Across both datasets, we tested 101 residue- and protein-level features at each experimental gate, plus 240 individual label probabilities from AstraSUIT2 and AstraROLE2. All tests used FDR correction within each gate. We report here only the strongest and most interpretable signals. We report controlled results for the CAR-T dataset (cysteine-free, K+E*<*30%) unless otherwise noted.

### CAR-T CD20 Benchmark: Multi-Gate Associations with Success

#### Expression Gate

Among the controlled subset (recovered *n*=3,019 vs. not recovered *n*=1,001), 53 features were significant (FDR*<*0.05). The strongest signals reflected topology-like sequence character: sequences with more inside-like character were more likely to be recovered (Table 1).

**Table 1:**
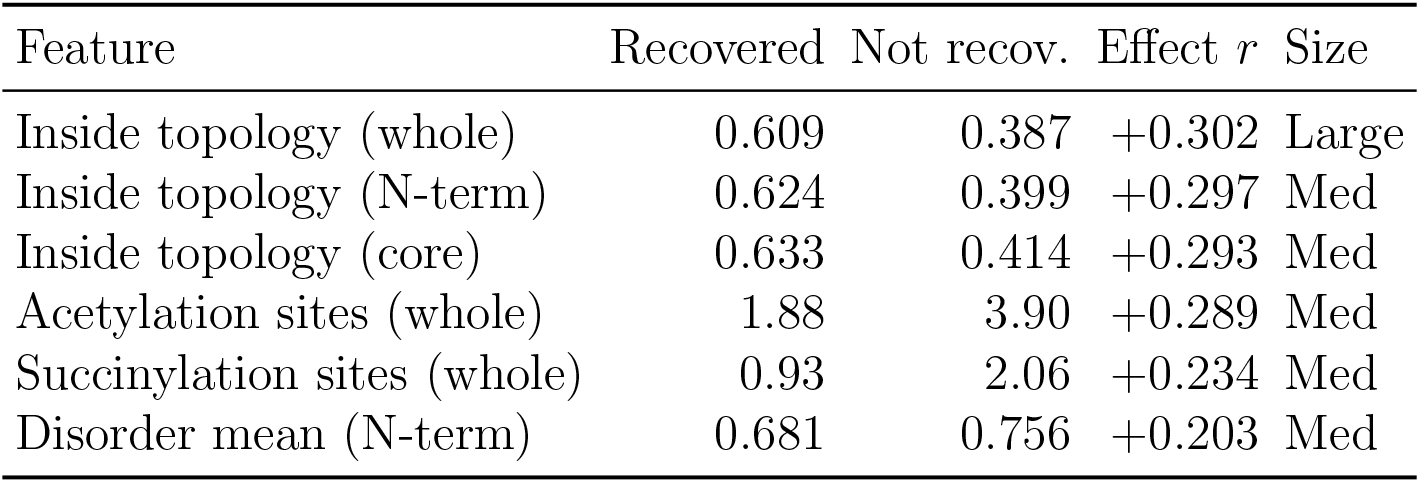
Top features associated with expression in the CAR-T benchmark (controlled).

| Feature | Recovered | Not recov. | Effect $r$ | Size |
| --- | --- | --- | --- | --- |
| Inside topology (whole) | 0.609 | 0.387 | +0.302 | Large |
| Inside topology (N-term) | 0.624 | 0.399 | +0.297 | Med |
| Inside topology (core) | 0.633 | 0.414 | +0.293 | Med |
| Acetylation sites (whole) | 1.88 | 3.90 | +0.289 | Med |
| Succinylation sites (whole) | 0.93 | 2.06 | +0.234 | Med |
| Disorder mean (N-term) | 0.681 | 0.756 | +0.203 | Med |

Recovered sequences were associated with substantially more inside-like topology character (Δ=+0.22, *r*=0.30) and fewer predicted acetylation and succinylation sites. Disorder was lower in recovered sequences across all regions, consistent with, but not establishing, a preference for compact, structured sequence profiles in T cell expression.

#### Enrichment Gate

Among enriched (*n*=418) vs. neutral (*n*=2,570) sequences in the controlled subset, 51 features were significant (Table 2).

**Table 2:** Top features associated with CD20-specific enrichment (controlled).

| Feature | Enriched | Neutral | Effect $r$ | Size |
| --- | --- | --- | --- | --- |
| PTM sites (whole) | 14.22 | 9.39 | +0.260 | Med |
| PTM sites (core) | 7.77 | 5.19 | +0.255 | Med |
| Phosphorylation sites (whole) | 5.88 | 4.23 | +0.221 | Med |
| Amyloid fraction (whole) | 0.537 | 0.754 | +0.218 | Med |
| Inside topology (core) | 0.479 | 0.658 | +0.193 | Med |
| Amyloid mean (whole) | 0.660 | 0.700 | +0.189 | Med |

Enriched binders were associated with significantly more predicted PTM sites (Δ=+4.8 sites), lower amyloid propensity (Δ= − 0.22 fraction), and less inside-like topology character. The amyloid-fraction signal was significant at the enrichment gate (*r*=0.22, *q<*10^−10^) but not at the expression gate (*r*=0.04, *q*=0.13), consistent with a gate-specific association at enrichment rather than expression. Additional label-level findings were examined but are not discussed here.

#### Depletion Gate

Among 166 depleted sequences (raw analysis; controlled *n*=31, too small), phosphorylation-related features showed the largest effects (Table 3). These findings are more tentative than the enrichment results due to the inability to control for known predictors.

**Table 3:**
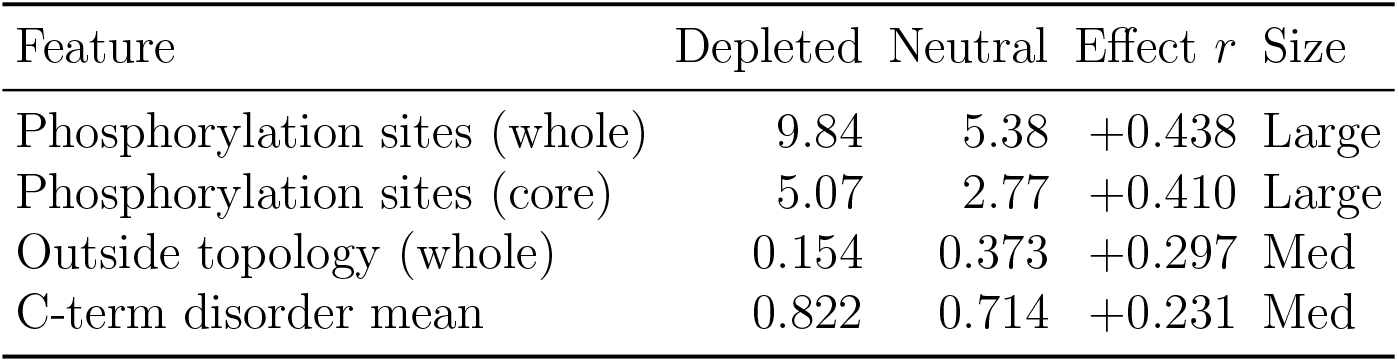
Top features associated with CD20-specific depletion (raw).

Depleted sequences had approximately twice as many predicted phosphorylation sites and lower outside-like topology character.

#### Expression–Enrichment Tradeoff

Several features flip direction between the expression and enrichment gates (Table 4). Properties associated with expression show the opposite direction of association at the enrichment gate.

**Table 4:** Features that flip direction between expression and enrichment gates (controlled).

| Feature | Expression $\Delta$ | Enrichment $\Delta$ | Direction |
| --- | --- | --- | --- |
| Inside topology (whole) | +0.221 | −0.165 | Opposite |
| PTM sites (whole) | −2.95 | +4.83 | Opposite |
| Acetylation sites (whole) | −2.02 | +1.16 | Opposite |
| C-term disorder mean | −0.008 | +0.069 | Opposite |

This pattern suggests that single-objective optimisation may favour sequences that perform differently across gates.

**Figure 3:**
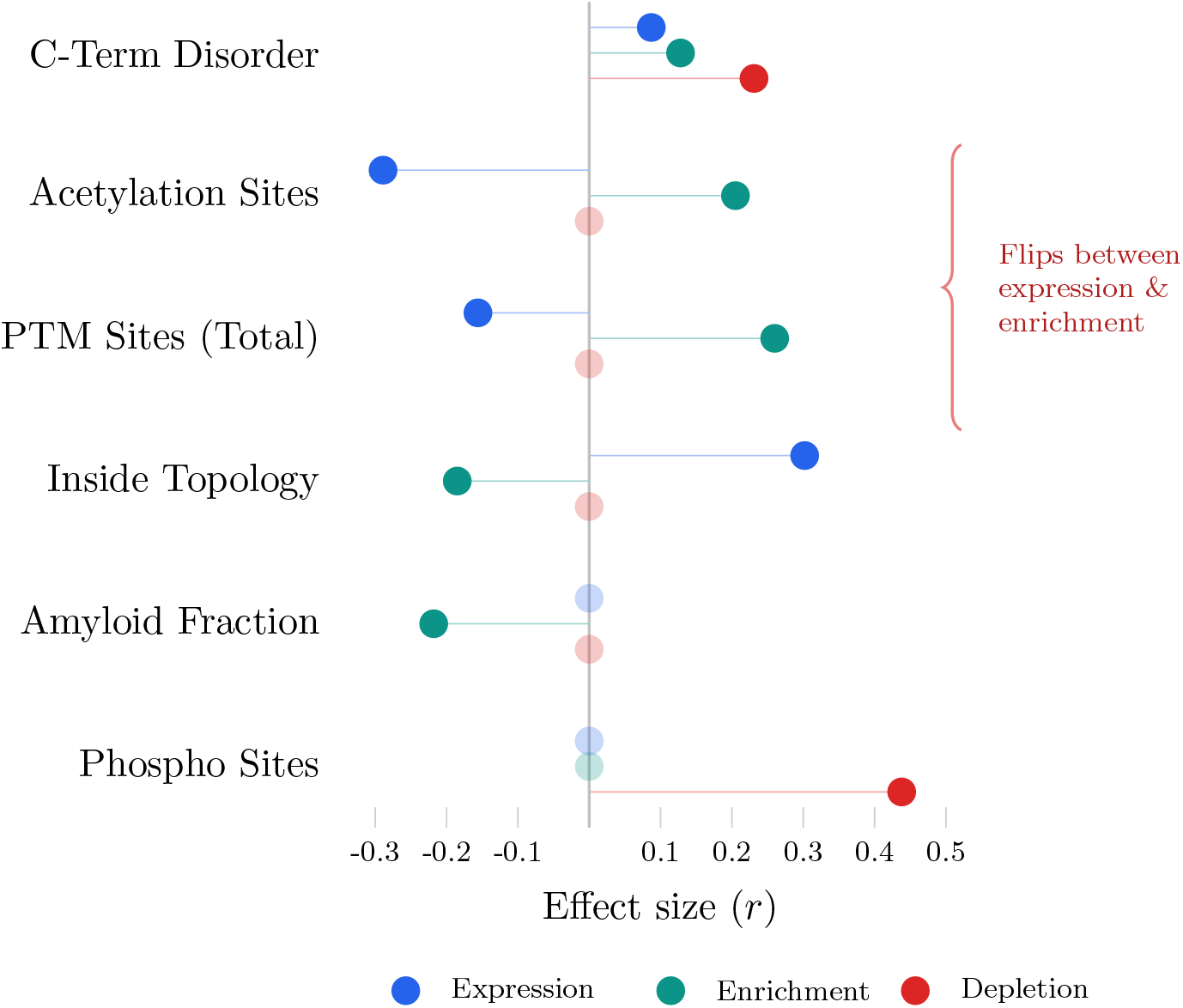
Multi-gate effect sizes in the CAR-T benchmark. The same sequence descriptor can carry different associations at expression, enrichment, and depletion gates, illustrating that biology-informed features may help flag which stage of the experimental pipeline is most at risk rather than providing a single overall score. Inside topology, PTM sites, and acetylation flip between expression and enrichment (brace). Phosphorylation is the dominant depletion signal. Faded points indicate non-significant features.

#### Filter Performance

We assessed the practical value of biology-informed features by applying them as sequential filters on the controlled subset (Table 5).

**Table 5:** Stacked filter performance on the CAR-T controlled subset.

| Filter stack | Enriched | Total | Hit rate | Lift |
| --- | --- | --- | --- | --- |
| Baseline (no-Cys, K+E<30%) | 418 | 3,019 | 13.8% | 1.0× |
| + not amyloidogenic | 194 | 830 | 23.4% | 1.7× |
| + outside topology >0.3 | 101 | 335 | 30.1% | 2.2× |
| + PTM sites $\geq 10$ | 71 | 184 | 38.6% | 2.8× |

Adding three biology-informed filters on top of the paper’s known heuristics raised the hit rate from 13.8% to 38.6%—a 2.8× lift—while simultaneously reducing the depletion rate from 1.0% to 0.5%. This filter analysis is a retrospective, within-benchmark exercise designed to illustrate the potential screening value of these features. The thresholds were selected post hoc on the same dataset used for evaluation; they should be treated as hypothesis-generating rather than as a validated predictive model. Prospective validation on independent data would be needed before these filters could be deployed in a production screening pipeline.

**Figure 4:**
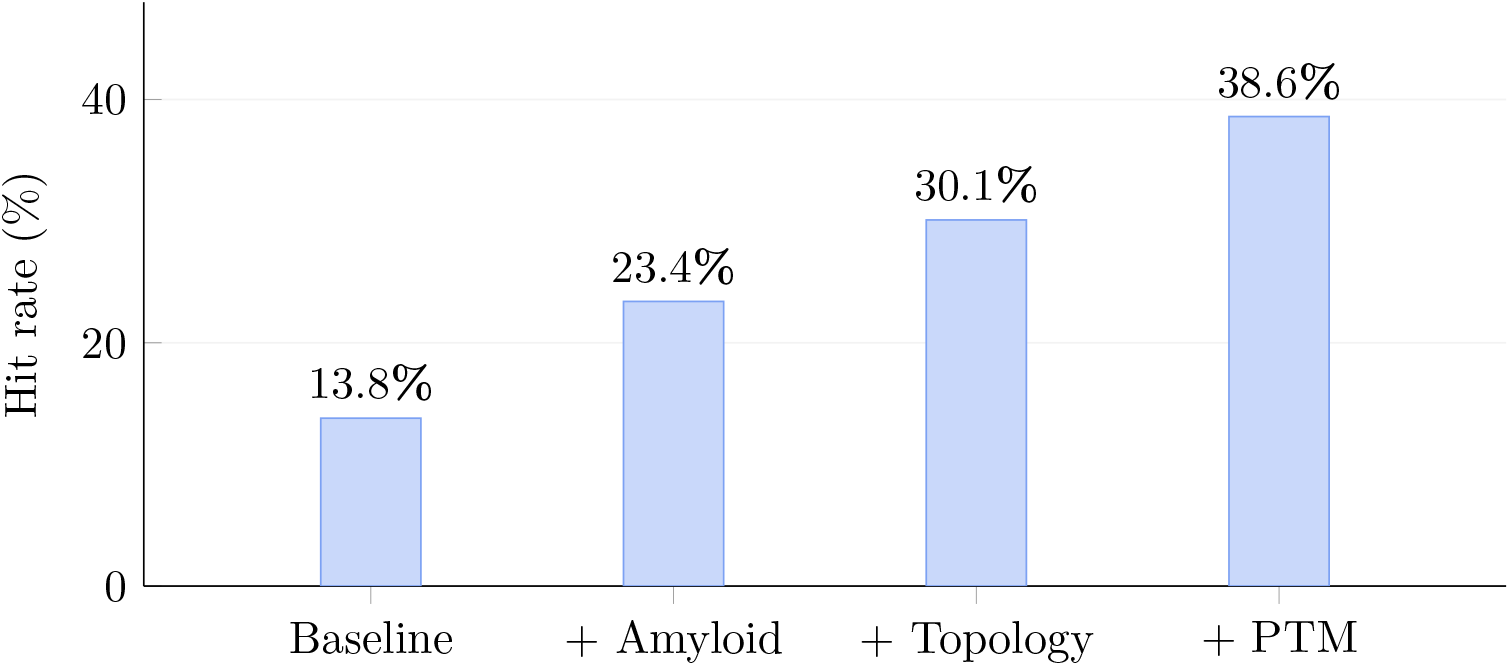
Stacked filter performance on the CAR-T benchmark. Each biology-informed filter is applied sequentially on the controlled subset (no-Cys, K+E*<*30% baseline). Hit rate increases from 13.8% to 38.6% (2.8*×* lift). Filters: (1) remove amyloidogenic sequences; (2) require outside topology *>*0.3; (3) require ≥10 predicted PTM sites.

#### EGFR Benchmark Tests Transferability in a Distinct Architecture and Assay Context

In the EGFR binding gate (62 binders vs. 541 non-binders), 26 features were significant (FDR*<*0.05). The dominant signals were markedly different from CAR-T (Table 6).

**Table 6:** Top features associated with binding in the EGFR benchmark.

| Feature | Binders | Non-binders | Effect $r$ | Size |
| --- | --- | --- | --- | --- |
| Disorder mean (whole) | 0.299 | 0.542 | +0.515 | Large |
| Disorder mean (C-term) | 0.364 | 0.628 | +0.484 | Large |
| Disorder fraction (whole) | 0.235 | 0.535 | +0.481 | Large |
| Disulfide sites (whole) | 8.27 | 3.60 | +0.440 | Large |
| Disulfide sites (core) | 4.46 | 1.74 | +0.439 | Large |
| Disorder fraction (C-term) | 0.318 | 0.662 | +0.414 | Large |

Low disorder was the dominant signal (*r*=0.52, large effect), with binders averaging 30% disorder probability versus 54% for non-binders. Predicted disulfide-related sequence character was also higher in binders (*r*=0.44), consistent with, though not proof of, a contribution from disulfide-stabilised architectures among successful designs. Lower amyloid propensity was again associated with binding success (*r*=0.28, *p*=3.9×10^−4^), replicating the CAR-T finding, and total predicted PTM sites were higher in binders (*r*=0.21, *p*=9.0×10^−3^). Additional label-level findings were examined but are not discussed here.

#### Length-confounding robustness

Because EGFR binder sequences span a wide length range (13–250 aa), count-based features (e.g., total PTM sites, disulfide sites) could be confounded by sequence length. Binders are on average longer than non-binders in this dataset. To assess robustness, we repeated the analysis with count features normalised by sequence length (sites per residue). The disorder and amyloid signals—which are already expressed as per-residue means or fractions—are inherently length-independent and remained significant. Normalised disulfide and PTM-total densities retained significance, though with attenuated effect sizes, confirming that the associations are not purely length-driven. Length remains a potential confounder in the EGFR analysis, and we note this among the study’s limitations.

The exploratory EGFR expression comparison (Round 2; *n*=22 non-expressed) is not interpreted further due to small sample size.

### Cross-Benchmark Synthesis Identifies Transferable and Context-Dependent Signals

#### Transferable Signals

Two feature families showed the same directional association with binding success in both datasets (Table 7).

**Table 7:** Transferable features: same direction, significant in both benchmarks.

| Feature | CAR-T $r$ | CAR-T $\Delta$ | EGFR $r$ | EGFR $\Delta$ |
| --- | --- | --- | --- | --- |
| Amyloid mean (whole) | +0.189 | −0.041 | +0.281 | −0.082 |
| PTM sites (whole) | +0.260 | +4.83 | +0.207 | +4.06 |
| PTM sites (core) | +0.255 | +2.59 | +0.265 | +2.41 |

Lower aggregation propensity was associated with binding success in both benchmarks. Higher predicted PTM-site density—which may reflect charged, surface-exposed sequence character rather than actual post-translational modification—was also associated with success in both univariate analyses, though in the heterogeneous EGFR dataset this signal is partly confounded by sequence length, making it a more robust independent signal in the fixed-length CAR-T context.

#### Architecture-Dependent Signals

Three feature families were significant in both datasets but with opposite directional associations (Table 8).

**Table 8:**
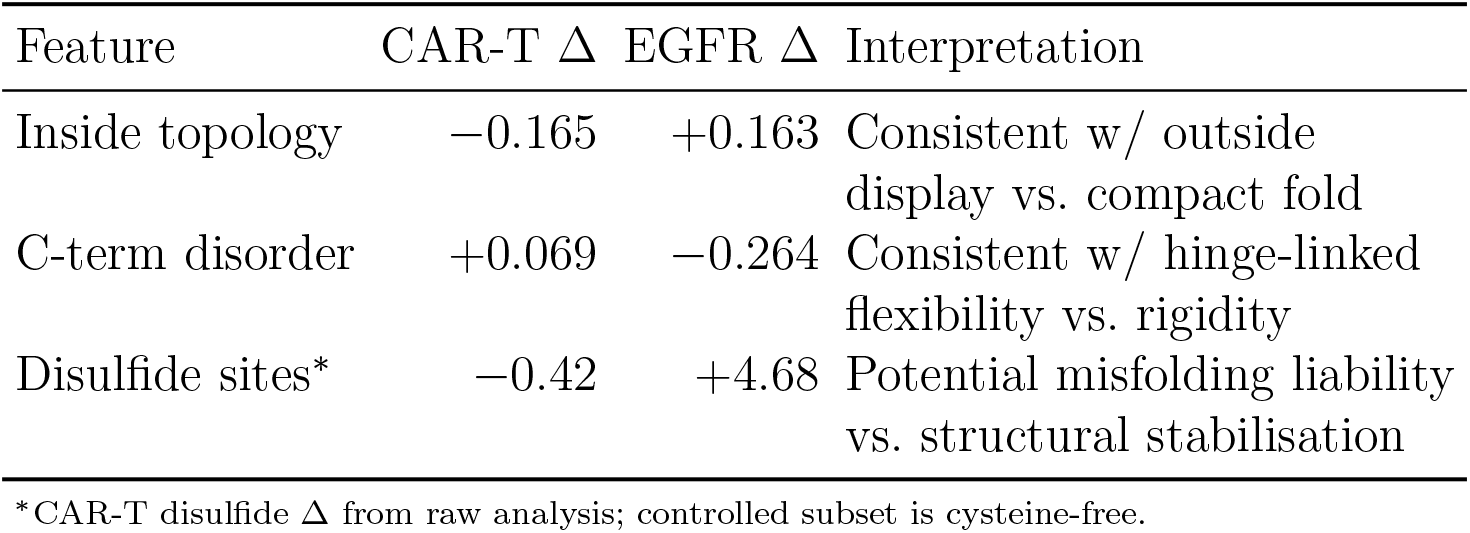
Architecture-dependent features: significant in both benchmarks, opposite direction.

| Feature | CAR-T $\Delta$ | EGFR $\Delta$ | Interpretation |
| --- | --- | --- | --- |
| Inside topology | -0.165 | +0.163 | Consistent w/ outside display vs. compact fold |
| C-term disorder | +0.069 | -0.264 | Consistent w/ hinge-linked flexibility vs. rigidity |
| Disulfide sites* | -0.42 | +4.68 | Potential misfolding liability vs. structural stabilisation |

\*CAR-T disulfide $\Delta$ from raw analysis; controlled subset is cysteine-free.

These features do not carry a fixed association with binder success. Their direction is consistent with the different requirements of each architectural context: membrane-displayed CAR domains may favour extracellular-like character and tolerate C-terminal flexibility, while standalone binders may favour compact, disulfide-stabilised folds. (The CAR-T disulfide Δ in Table 8 is drawn from the raw analysis because the controlled subset excludes cysteine-containing sequences, in which disulfide signal resides.)

**Figure 5:**
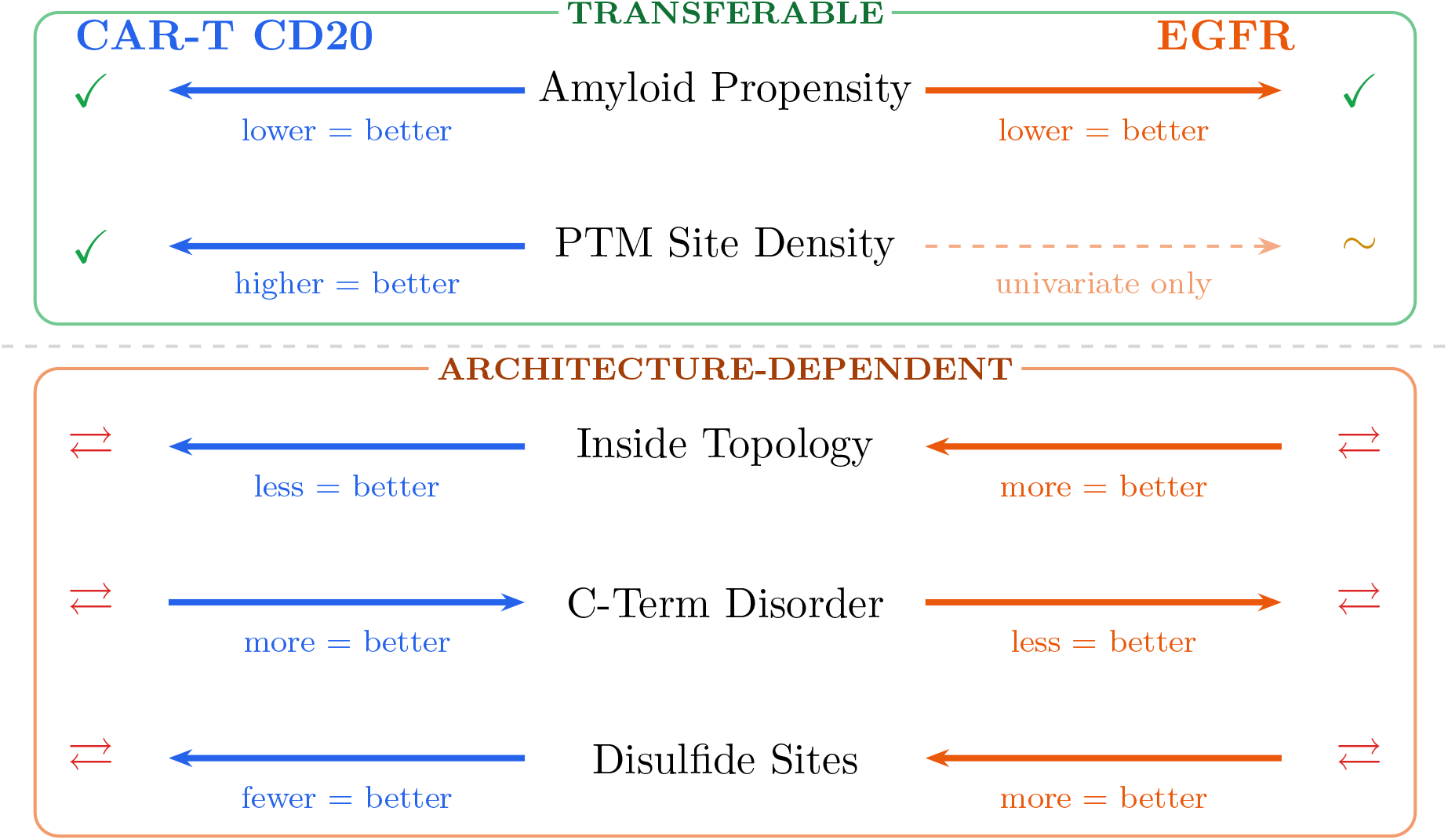
Cross-benchmark signal comparison. Amyloid propensity (top) shows the same directional association with binding success in both benchmarks. PTM-site density shows the same univariate direction but is partly confounded by sequence length in the heterogeneous EGFR dataset (dashed arrow). Architecture-dependent descriptors (bottom, orange border) flip direction: properties associated with success in CAR-T are associated with failure in EGFR, and vice versa, consistent with their different structural requirements.

**Figure 6:**
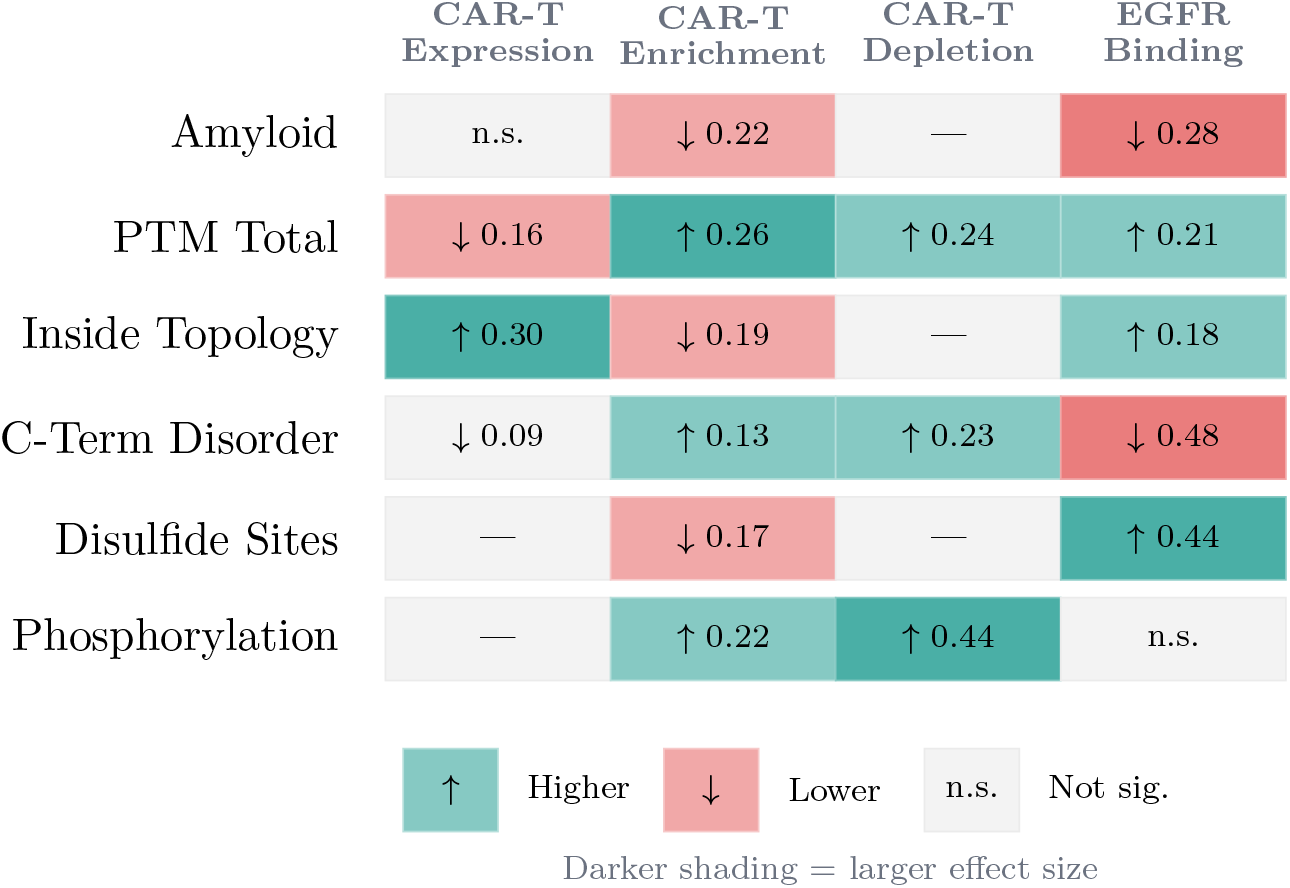
Summary of effect directions across gates and benchmarks. Each cell shows the direction of association (arrow) and effect size (*r*) for the success group at each gate. Teal cells indicate higher values in the success group; red cells indicate lower values. Darker shading indicates larger effect sizes. Grey cells indicate non-significant or untested comparisons. CAR-T values are from the controlled subset except for the depletion column (raw; controlled *n* too small) and the disulfide row (raw; controlled subset is cysteine-free).

#### Context-Specific Signals

Several signals were significant in only one dataset:

- **CAR-T phosphorylation–depletion signal**: predicted phosphorylation sites showed a notable association with T cell depletion (*r*=0.44, large; raw analysis only, see Limitations), with no equivalent in the EGFR dataset.
- **CAR-T expression–enrichment tradeoff** : inside-like topology, PTM burden, and acetylation sites all flipped direction between expression and enrichment gates—a pattern unique to the multi-gate CAR-T assay.
- **EGFR disorder dominance**: low disorder was the single strongest signal for EGFR binding (*r*=0.52), far exceeding any single CAR-T feature effect size. This is consistent with the different structural requirements of standalone binders.
- **Receptor-category context-dependence**: the receptor protein category label was associated with binding success in EGFR (*r*=0.32) but with depletion in CAR-T (*r*=0.33), illustrating that the same model output can carry opposite associations across design contexts.

### EGFR Scaffold-Stratified Analysis

Scaffold class (assigned by length: peptide *<*50 aa, miniprotein 50–90, nanobody 91–150, scFv *>*150) was significantly associated with EGFR binding (*χ*^2^=10.6, *p*=0.014). To test whether the key signals are genuine within-class effects, we repeated univariate tests within each class with ≥10 binders (Table 9).

**Table 9:** Within-scaffold-class signal consistency in EGFR binding.

| Feature | Miniprotein<br>($n=23$ bind.) | Nanobody<br>($n=12$ bind.) | scFv<br>($n=20$ bind.) |
| --- | --- | --- | --- |
| Disorder | ↓ sig. | ↓ n.s. | ↓ sig. |
| Disulfide | ↑ sig. | ↑ n.s. | ↑ sig. |
| Amyloid | ↓ sig. | ↓ n.s. | ↓ sig. |
| PTM total | ↑ n.s. | ↑ n.s. | ↑ sig. |

All four features maintained the same direction within every scaffold class (zero reversals across 12 comparisons). Disorder, disulfide, and amyloid reached significance within both miniprotein and scFv classes independently. PTM-total was the weakest within-class signal. The core EGFR signals reflect genuine sequence-level properties, not scaffold composition artefacts.

## Framework for Context-Aware Pre-Synthesis Screening

Based on the retrospective associations observed across both benchmarks, we outline a candidate layered screening approach for de novo binder design. This framework is hypothesis-generating: the filter thresholds and layer assignments are derived from the same data used to identify the signals, and prospective validation would be required before adoption.

### Universal Filters

- **Aggregation risk**: may help deprioritise sequences with high predicted amyloid propensity (CAR-T: 1.7*×* lift; EGFR: 1.3*×* lift in retrospective analysis).
- **PTM-site density**: may be worth favouring sequences with higher predicted PTM-site counts (independently informative in CAR-T; partly confounded by length in EGFR).

### Architecture-Aware Filters

Three feature families may require direction to be adapted to the binder format:

- **Topology-like character**: outside-like may be preferable for membrane-displayed binders; inside-like for standalone binders.
- **Disorder**: moderate flexibility may suit CAR-type binders; low disorder may suit standalone binders.
- **Disulfide potential**: may favour standalone architectures; may act as a risk indicator in multi-domain constructs.

### Context-Specific Filters

- CAR-T: phosphorylation-site density and receptor-like character as tentative depletion risk indicators; consider balancing expression and enrichment gates.
- Standalone binders: disorder stringency may warrant heavier weighting.

### Transfer and Non-Transfer

The CAR-T filter stack (amyloid + outside topology + PTM ≥ 10) achieves a 2.8× lift within the CAR-T benchmark. Only amyloid transfers robustly to EGFR; PTM-total transfers partially; topology actively *hurts* standalone binders (0.8×). The proposed three-layer workflow (Figure 7) adapts screening to binder architecture.

**Figure 7:**
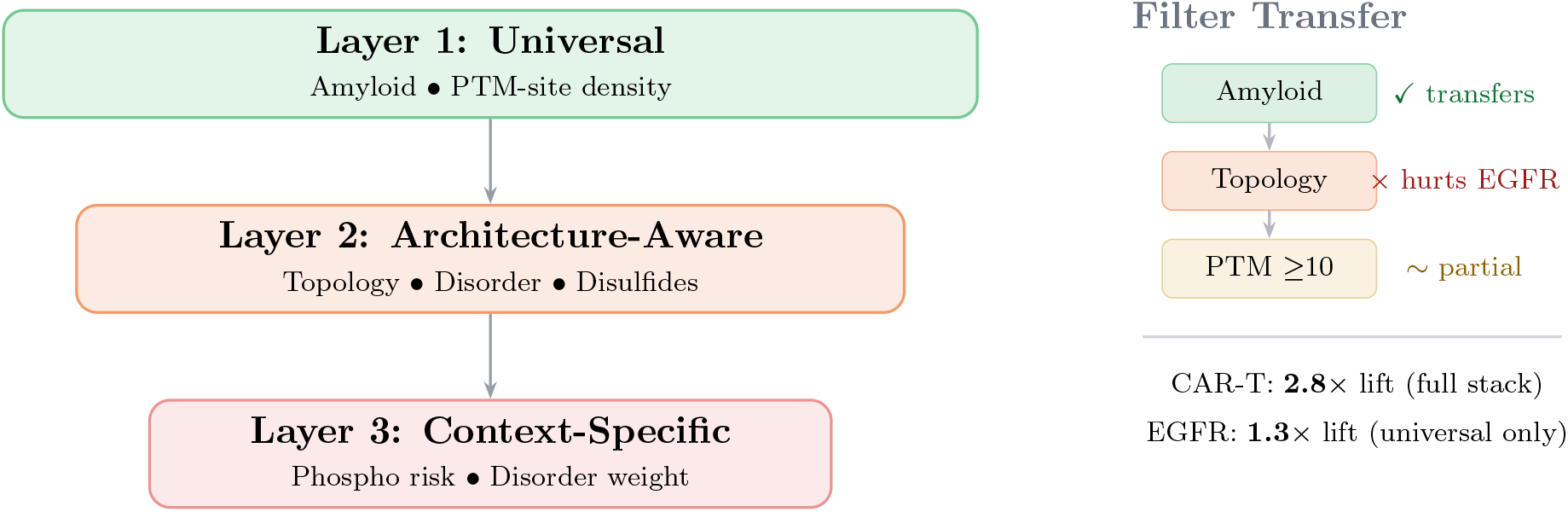
Context-aware screening framework and filter transfer. Left: proposed three-layer workflow. Right: the CAR-T filter stack achieves 2.8× lift, but only universal filters transfer to EGFR; topology hurts standalone binders, illustrating that screening should be adapted to binder architecture.

## Discussion

### Biology-Informed Features as a Complementary Screening Layer

The original benchmark papers [1, 2] established that simple sequence-level heuristics outperform complex structural scoring functions, but did not explore biology-informed ML features. The present analysis identifies specific feature families—amyloidogenicity, disorder, topology-like character, disulfide-related signals, and (in the fixed-length CAR-T setting) PTM-site density—that carry medium-to-large effect-size associations with experimental outcomes after controlling for the known heuristics, provides a cross-benchmark comparison of which associations transfer, and proposes a layered screening framework.

Biology-informed descriptors are most likely to add value in settings where simple composition rules are insufficient—diverse scaffolds, variable-length designs, or novel architectural contexts.

Beyond broad success/failure ranking, this study suggests three practical uses for biology-informed sequence descriptors. First, they can help indicate which experimental gate may be most at risk—for example, expression versus enrichment in CAR-T. Second, they can add complementary signal beyond structure-confidence metrics, particularly in heterogeneous binder settings. Third, they can show when the same descriptor carries opposite associations across deployment contexts, implying that screening rules should be architecture-aware rather than universal.

### Why the Same Signal Can Flip Across Design Contexts

A central observation from this study is that biology-informed sequence descriptors do not carry fixed associations across binder-design tasks; their interpretation depends on how a binder is deployed. Several descriptors show opposite associations with success depending on the binder architecture, without implying a single mechanistic explanation.

Consider topology-like sequence character. In the CAR-T benchmark, sequences with more outside-like character were more likely to show CD20-specific enrichment. This is consistent with the binder’s role as an extracellular domain displayed on a membrane receptor: outside-like sequence character may reflect compatibility with extracellular display and target engagement across an immunological synapse. In the EGFR benchmark, by contrast, successful binders were enriched for more inside-like (compact, globular) character, consistent with the need for standalone binders—nanobodies, miniproteins, scFvs—to fold into stable, self-contained structures with hydrophobic cores.

A similar pattern holds for disorder. The positive association between C-terminal disorder and CAR-T enrichment is consistent with hinge-linked flexibility requirements [11], while EGFR binders show the opposite—low disorder is associated with success, consistent with the need for rigid, well-ordered folds in standalone architectures. And for disulfide potential: associated with success in standalone antibody-derived architectures where disulfides stabilise the fold, but with failure in the multi-domain CAR construct context, where they may represent a potential misfolding liability.

These flips are unlikely to reflect simple lack of replication and are consistent with differing architectural requirements. They suggest that the same descriptor output should be interpreted differently depending on the architectural context, and that the underlying mechanisms linking these descriptors to experimental outcomes remain open questions for future investigation.

### Limitations and Interpretation Boundaries

Several limitations should be noted.

First, this is a retrospective re-analysis of published benchmark data. We did not design, synthesise, or test any binders; all associations are correlational.

Second, we did not experimentally validate the proposed mechanisms. For example, the association between amyloidogenicity and binder failure is consistent with the known role of scFv clustering and aggregation in driving antigen-independent tonic signalling in CAR-T cells [10], and the topology and disorder findings are consistent with known sensitivity to hinge and spacer design [11], but these remain interpretive hypotheses rather than confirmed mechanisms.

Third, the models were trained on natural proteins and applied to synthetic sequences; their outputs are sequence compatibility indicators, not confirmed biochemistry (as detailed in Methods).

Fourth, the EGFR dataset is smaller (*n*=603) and heterogeneous in binder type and length. Count-based features may be confounded by length; normalised analyses retained significance with attenuated effect sizes.

Fifth, certain gates have limited sample sizes (CAR-T depletion controlled *n*=31; EGFR strong-binding *n*=30; EGFR expression *n*=22 non-expressed). The depletion-gate findings are among the more tentative results—the controlled subset was too small for independent confirmation, and the raw analysis does not control for known sequence-level predictors.

### Future Directions

Prospective validation—designing binders with and without biology-informed filtering and comparing experimental outcomes—would provide the strongest test of these findings. Additional public benchmarks (e.g., future rounds of Bits to Binders or Adaptyv competitions, or the Protein Engineering Tournament) could further test transferability. Targeted wet-lab experiments could validate specific failure-mode hypotheses, such as whether predicted amyloidogenicity correlates with surface clustering or tonic signaling markers. Finally, integrating context-aware biology-informed features into generative design pipelines—as constraints during sequence generation rather than post-hoc filters— represents a natural next step.

**Figure 8:**
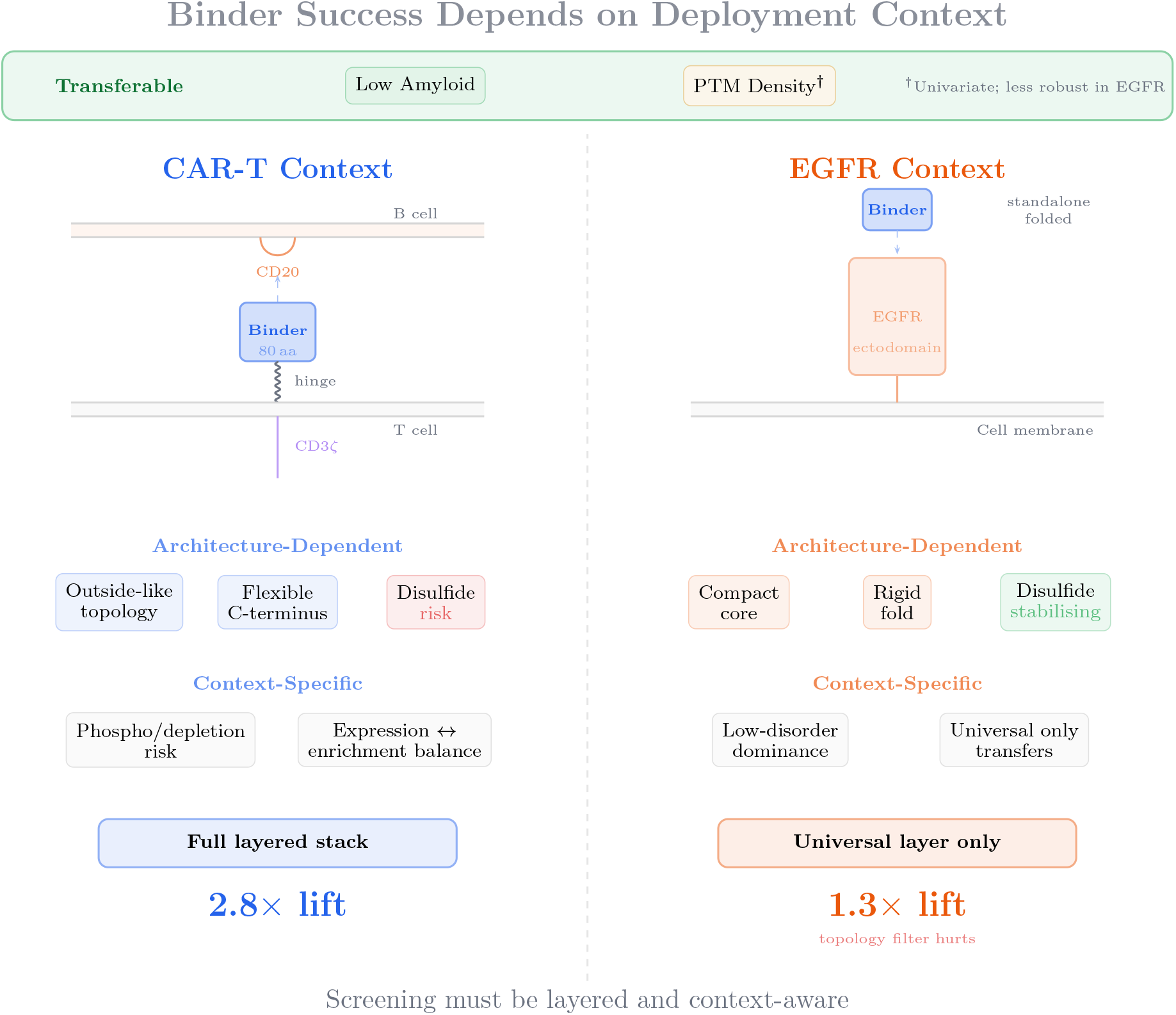
Binder success depends on deployment context. Low amyloid propensity is the most robust transferable signal; PTM-site density is a recurring univariate correlate but partly confounded by length in EGFR (dagger). Architecture-dependent descriptors (topology, disorder, disulfides) carry opposite associations in membrane-displayed CAR-T binders versus standalone EGFR binders. Context-specific factors (phosphorylation/depletion association, expression–enrichment balance) are tied to individual deployment settings. The full layered filter stack achieves 2.8× lift in CAR-T; only the universal layer transfers to EGFR.

## Conclusion

Re-analysis of two public binder-design benchmarks with biology-informed ML-derived sequence features identifies three layers of signal:

- **Aggregation propensity** is the most robust transferable signal. **PTM-site density** shows a univariate association in both benchmarks but is partly confounded by length in EGFR.
- **Topology, disorder, and disulfide character** flip direction between contexts— consistent with the different requirements of membrane-anchored versus standalone architectures.
- The **expression–enrichment tradeoff** and a tentative **phosphorylation– depletion association** are specific to CAR-T; **low disorder dominance** is specific to standalone binders.

These findings suggest that binder evaluation may benefit from moving beyond structure- and affinity-centred scoring toward context-aware, multi-gate screening that accounts for expression compatibility, aggregation risk, and architectural fit. In practice, these descriptors may be most useful when screening needs to distinguish not just whether a candidate looks risky, but whether the risk is more consistent with expression failure, aggregation burden, architecture mismatch, or loss of functional compatibility in a specific deployment setting.

## Data and Code Availability

All binder sequences and their corresponding experimental outcomes for the CAR-T CD20 benchmark are publicly accessible via the Bits to Binders data package [1]. EGFR binder sequences and binding results are available via the Adaptyv Bio competition repositories [2, 8].

Computational predictions were generated using Orbion’s Astra ML model suite (AstraUNFOLD, AstraPTM2, AstraSUIT2, AstraROLE2). The suite is commercially available via the Orbion platform; to support open-science initiatives, access is provided free of charge to academic researchers through the web interface at https://app.orbion.life/. Results presented in this study can be replicated using the publicly available sequences and the Orbion platform.

### Model interpretability caveat

While the Astra ML suite functioned as an effective source of biology-informed sequence features in this study, the proprietary nature of the training datasets and underlying model weights limits complete mechanistic interpretability of the computational predictions. The associations identified here should be understood as statistically validated patterns correlated with experimental outcomes, not as direct proofs of molecular mechanism. Future work should seek to cross-validate these findings using orthogonal structural methods (crystallography, cryo-EM, molecular dynamics) and open-source prediction tools.

